# Inferring single-cell gene expression profiles from multicellular spatial transcriptomics data with st2sc

**DOI:** 10.64898/2026.11.10.765653

**Authors:** Young Je Lee, Swayam Shah, Shaurya Mantrala, Jose Lugo-Martinez

## Abstract

Spatial transcriptomics (ST) enables the quantitative measurement of gene expression within tissue sections but is currently limited in spatial resolution, as each spot aggregates signals from multiple cells. Recovering single-cell gene expression profiles from these multicellular measurements requires two steps: explicit identification of individual cells and accurate estimation of per-gene expression. While existing methods have demonstrated the feasibility of recovering single-cell expression profiles, accurately estimating gene-level expression for individual cells remains an important challenge. To address this challenge, we propose st2sc (Spatial Transcriptomics to Single-Cell), an end-to-end framework for inferring single-cell gene profiles from multicellular ST data. st2sc introduces a novel Attention Gradient Flow Tracking (AGFT) algorithm that extracts attention maps from a Vision-Gene Transformer to identify candidate cell instances without predefined background constraints. The detected instances are then used by a set of regressors trained under a multiple-instance, multiple-target regression setting to predict precise single-cell gene expression counts. Comprehensive evaluations on synthetic and Visium-like datasets demonstrate that st2sc significantly outperforms state-of-the-art methods in accurately segmenting cellular instances and predicting gene counts. The method can further resolve contested 2um bin in Visium HD. Application to human IPF lung showed the model can recover the subtype marker level analysis. This advancement further improves single-cell resolution analysis in spatial transcriptomics, facilitating more accurate downstream biological analyses.

## 1 Introduction

The field of spatially resolved transcriptomics (ST) is rapidly bridging the gap between whole-transcriptome gene expression profiling and the rich morphological context provided by tissue imaging. By jointly measuring spatial location and transcriptional activity, ST has transformed our ability to study cellular organization, tissue architecture, and microenvironment-driven gene regulation. While early ST technologies successfully mapped gene activity across tissue sections, the persistent challenge has been to achieve true single-cell resolution while maintaining unbiased coverage of the entire transcriptome. This pursuit has driven intense innovation, with technologies such as Slide-seqV2 [1] and Stereo-seq [2] achieving near single-cell resolution (≈10*µm*), and the recent commercial introduction of Visium HD [3] further increasing precision with 2×2*µm* barcoded squares.

Despite these technological advances, widely used ST methods still operate at near single-cell or multicellular resolution, leaving a critical need for computational tools to infer single-cell expression profiles to facilitate more accurate downstream biological insights. Several computational methods have demonstrated the feasibility of deconvoluting multicellular ST data, including SpatialScope [4] and scResolve [5]. By computationally integrating high-resolution histology with spatially resolved gene expression data and single-cell RNA sequencing (scRNA-seq) reference profiles, these tools enable the deconvolution of transcriptomes at single-cell resolution, thereby elucidating functional intra-type heterogeneity and detecting rare cell populations within the tissue microenvironment. However, these existing computational approaches face significant limitations. They often rely on the relative accuracy of cell-type prediction rather than the absolute accuracy of inferred gene counts. Crucially, they suffer from high computational cost and heavy dependency on method-specific parameters (e.g., metagenes in scResolve) or depend on the availability and quality of matched external scRNA-seq references (in SpatialScope). Furthermore, their reliance on nuclei segmentation often disregards informative histological features and limits their ability to accurately model full cell structure. Importantly, increasing spatial resolution does not entirely eliminate ambiguity in assigning expression counts to individual cells. Even at the 2*µm* resolution of Visium HD, a single expression bin may span a pair of neighboring cells, similar to how a Visium spot captures transcripts from several cells. Analysis methods will either assign all expression counts to the one cell (bin2cell [6]) or use cluster-level information to assign the expression counts (ENACT [7]), but the first discards the second cell’s share entirely, and the latter may divides the counts evenly whenever both cells fall in the same cluster.

To address these limitations, we propose st2sc (Spatial Transcriptomics to Single-Cell), an end-to-end framework for inferring high-accuracy single-cell gene profiles from multicellular spatial transcriptomics data. Our approach bypasses the dependencies and computational bottlenecks of prior work by combining a novel segmentation algorithm, inspired by self-supervised learning [8] and gradient flow tracking [9], with a multiple-instance, multiple-target regression framework. This architecture is designed to simultaneously identify individual cell instances directly from histology and recover their true absolute gene expression profiles, marking a significant step toward achieving robust end-to-end single-cell resolution in spatial transcriptomics. Through both synthetic and real-world simulations, st2sc demonstrates more accurate cell localization and achieves substantially improved absolute gene count recovery compared to existing methods. We further evaluated st2sc on Visium HD data, demonstrating its ability to resolve contested 2 µm expression bins by improving the assignment of cell-type-specific marker gene counts compared with existing methods. To demonstrate its biological applicability, we applied st2sc to human idiopathic pulmonary fibrosis (IPF) lung Visium data, recovering fibroblast expression gradients near fibrotic foci and spatial accumulation patterns of SPP1+ macrophages consistent with findings from higher-resolution spatial transcriptomics. These findings provide strong evidence that st2sc provides an improved framework for analyzing complex multicellular spatial transcriptomics data, enabling the extraction of enhanced accuracy and depth of biological insights compared to existing methodologies. st2sc is available on GitHub: https://github.com/youngje-cs-cmu/st2sc

## 2 Method

In this section, we present st2sc, a multiple-instance multiple-target regression framework designed to reconstruct single-cell expression profiles from multicellular resolution spatial transcriptomics. We begin by formally framing the biological problem as a multiple-instance, multiple-target regression problem and outlining the overall structure of the proposed framework. We then introduce the development of our instance identification Attention Gradient Flow Tracking (AGFT) algorithm. Finally, we describe the dataset construction and evaluation methodology.

### 2.1 Problem formulation

Our primary goal is to recover single-cell expression from multicellular spatial transcriptomics. To solve this biological problem, let us first formally define the multiple-instance, multiple-target regression problem. Let *N* denote the number of bags (spots) and *M* the number of targets (genes). We index the spots by *i* ∈ {1, … , *N*} and the targets by *j* ∈ {1, … , *M*}. Each bag *i* contains a set of instances, denoted by 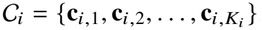, where *K*_*i*_ is the number of instances (cells) in the spot *i*. Here, **c**_*i*,*k*_ ∈ ℝ^*M*^ is the gene expression vector for the *k*-th cell in spot *i*. The goal of multiple-instance, multiple-target regression is to learn a function *f* that maps the set of instance vectors C_*i*_ to the observed spot-level target vector **y**_*i*_ = [*y*_*i*,1_, *y*_*i*,2_, … , *y*_*i*,*M*_]^*T*^. We aim to find *f* such that *f* (C_*i*_) ≈ **y**_*i*_ for all *i* = 1, … , *N*.

In the context of our problem of recovering true single-cell expression from multicellular spatial transcriptomics, each bag corresponds to a spot *i* ∈ {1, … , *N*}, and target values are gene expressions of *M* genes. For each cell **c**_*i*,*k*_ within the spot, our aim is to regress the *M* genes. In this setting, we assume that the observed spot-level expression is an aggregation of the underlying cellular expressions, expressed as:

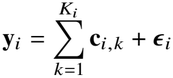

where *K*_*i*_ is the number of cells at the spot *i*, **c**_*i*,*k*_ is the true single-cell gene expression vector and *ϵ* _*i*_ represents noise. We are also given a histology image *I* with width *w* and height *h*, from which we can extract the image patch *I*_*i*_ corresponding to the spot *i* and another image patch 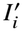 which is larger image than *I*_*i*_ that can provide more visual context for the model.

One instance of this problem is Visium HD. Visium HD measures gene expression using 2*µ*m × 2*µ*m spatial bins, providing substantially higher spatial resolution than Visium, its predecessor. However, a single bin may overlap a pair of neighboring cells, making it difficult to attribute the observed gene counts to individual cells. This creates a multicellular resolution problem even in near-single-cell-resolution data. Importantly, bins overlapping only one identified cell provide an opportunity for model training, under the assumption that their measured transcripts originate from that cell. Using these bins, a model can learn the relationship between cellular image features and gene expression, which can subsequently be applied to contested bins overlapping multiple cells.

### 2.2 Overview of st2sc framework

Our problem formulation considers solving a multiple-instance, multiple-target regression problem to recover single-cell gene expression, which introduces an additional challenge of identification of the instances that are expected to correspond to individual cells. To address this challenge, we propose the **st2sc** framework, an end-to-end framework to recover the true expression, which is shown in Figure 1A. The framework consists of two modules: (1) *Attention gradient flow tracking algorithm*, which identifies instances (cells) within each spot (*cell identification stage*), and (2) *Regressors* that solves the multiple-instance, multiple-target regression problem using the identified instances and the corresponding histology image (*gene expression inference stage*). Formally, for each spot *N*_*i*_, we first detect the set of instances from its corresponding image patch *I*_*i*_ using the attention gradient flow tracking algorithm. Given these identified instances, the regressors then predict the target gene expression vector 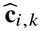 for each cell *k* by solving the multiple-instance multiple-target regression problem. These cell-level predictions are subsequently aggregated to reconstruct the observed spot-level expression, providing weak supervision for model training. In the following paragraphs, we describe each component in detail.

**Fig. 1.**
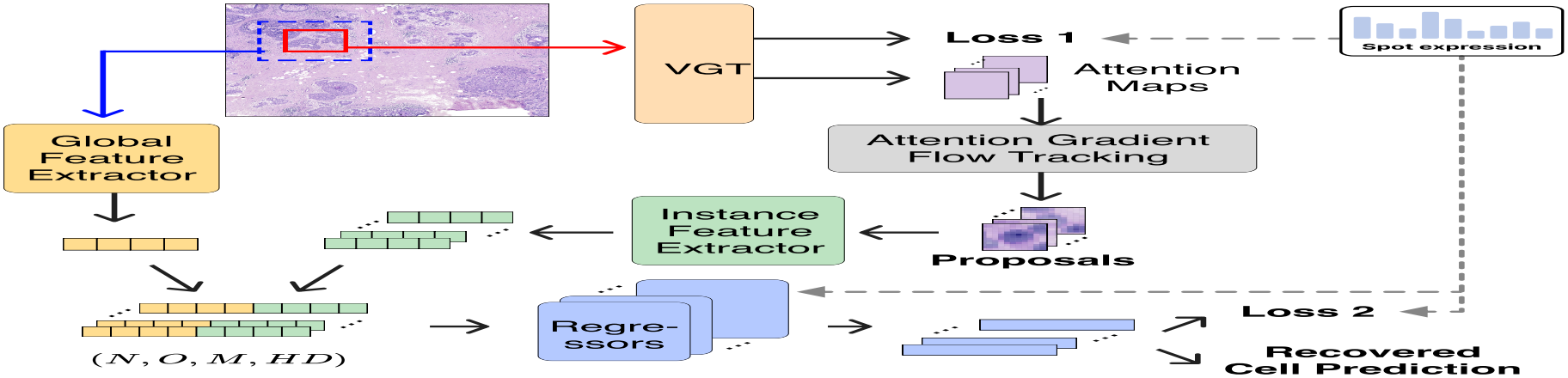
Overview of st2sc. Schematic overview of st2sc. The st2sc framework is designed to recover single-cell expression profiles from multicellular spatial transcriptomics data using a multiple-instance, multiple-target regression approach. The method first identifies and generates cell instance proposals from spatial spots using a Vision-Gene Transformer (VGT) combined with attention gradient flow tracking. Next, st2sc performs regression using an ensemble of small neural networks to produce a cell-by-gene expression matrix, thereby reconstructing the single-cell profiles.

#### Vision-Gene transformer (VGT)

First, we describe the Vision-Gene transformer encoder used to extract attention maps. First, we revisit the concept of the multi-head self-attention mechanism. Let *q* be a query, *k* a key, and *v* a value. Given an input sequence *s*, the self-attention of the input *S A*(*s*) can be computed as:

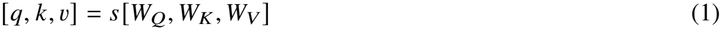

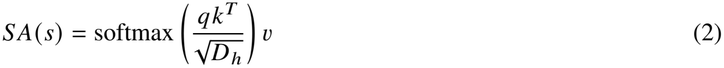

where *h*_*Q*_ *(s)* , *h*_*K*_ *(s)* , *h*_*V*_ *(s)* are linear projection layers and *W*_*Q*_, *W*_*K*_ , *W*_*V*_ are the corresponding learnable weight matrices. We initialize the Vision-Gene Transformer, illustrated in (Supp. Fig. 1A). The VGT architecture follows that of a Vision Transformer (ViT), where the input image is divided into patches that are linearly projected into an embedding space. However, the Vision-Gene Transformer differs from the standard ViT in two key aspects: (1) no positional encoding is used, as it adds noise in the attention gradient flow tracking, and (2) a gene encoder replaces the class token. Specifically, *M* gene embeddings are initialized and concatenated with the visual patch embeddings before being processed by the transformer encoder. As a result, the input sequence *s* to the Transformer encoder layer in the Vision-Gene Transformer becomes the concatenation of linear embedding from spot image *I*_*i*_ and gene embedding. The output corresponding to the gene encoder is subsequently mapped to the spot-level gene expression. This design can be interpreted as an early fusion of visual and transcriptomic features, allowing the model to learn a joint representation that captures morphological cues relevant to gene expression. Therefore, we expect the Vision-Gene Transformer to attend to informative histological regions conditioned on observed spot-level expression.

#### Attention gradient flow tracking

Next, we describe the attention gradient flow tracking algorithm. Conceptually, it extends the gradient flow tracking method introduced by Li et al. [9], but adapts it to a probabilistic attention map rather than a discrete image. In our setting, the input is a continuous-valued, image-like map in probabilistic space, without a predefined background region (Figure 2B). The algorithm proceeds as follows: From the Vision-Gene Transformer, we extract all attention maps. Because the visual and gene encoders share the same attention distribution due to early fusion, we first isolate the visual component. The attention maps are then resized to the resolution of the image patch, forming an image-like representation *A*^′^. For computational efficiency, we take the average of all extracted attention maps *A*_*avg*_. The gradient flow tracking algorithm is then applied to *A*_*avg*_, evaluating the gradient flow for all pixels, rather than relying on background thresholding as in traditional image-based methods [9, 10]. This design prevents performance bias introduced by threshold selection. The gradient flow of each pixel is then traced until the turning angle becomes less than or equal to zero, at which point the endpoint is marked as a sink and the trajectory is recorded. The resulting flow sinks correspond to the predicted cell instance centers, which are subsequently refined during postprocessing.

**Fig. 2.**
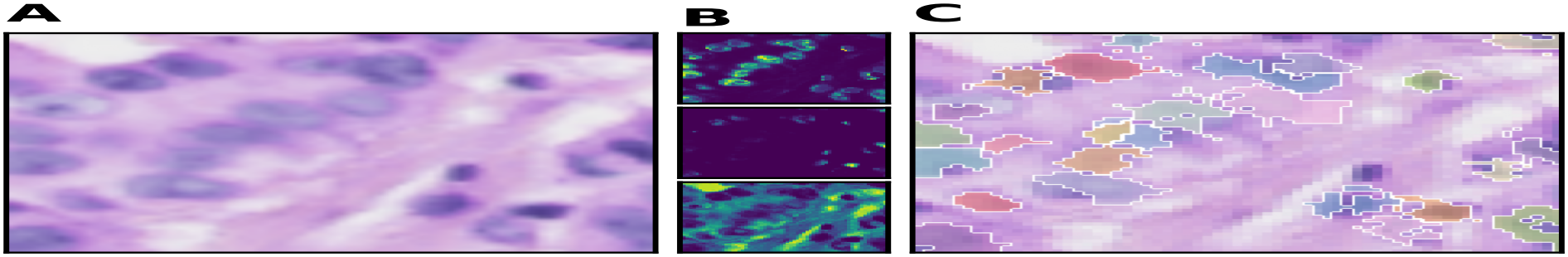
Attention gradient flow tracking (AGFT) captures cell as instance. Although attention distribution varies on instances across different heads and genes, the Attention gradient flow tracking captures the cells, providing instance proposals. (A) An input histology image (B) Attention maps extracted from multi-heads. Brighter (yellow) pixels indicate higher attention, darker (blue) pixels indicate lower attention. (C) Ouptut of the AGFT algorithm after postprocessing, where each color represents gradient flows exited in the same sink.

#### Regressors

Lastly, from the attention gradient flow tracking algorithm, we obtain a set of instance proposals in the form of masks. In this step, we train the second phase of the framework, **regressors**, to solve the multiple-instance, multiple-target regression problem. The central idea is to train a lightweight model for each gene rather than a single large model, thereby accommodating heteroscedasticity across genes. For each spot *i*, the **regressors** phase takes two inputs: (1) a group of instance proposals mask (*O, r*_*x*_, *r*_*y*_), and (2) the larger patch image 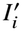. Let *f*(·) be a neural network. We define four functional components: (1) a visual feature extractor of the larger patch *f*_*PatchImg*_ (·), (2) a visual feature extractor of instances *f*_*InstanceImg*_ (·), a non-learnable pooling operator over the output of *f*_*PatchImg*_ (·), (3) an *aggregator f*_*Aggregate*_ *x* , a non-learnable aggregator function defined as *y* = *Wx*+*b, W* = **1**^⊤^ ∈ ℝ^1×*O*^, *b* = 0. Finally, we define (4) a group of regressors

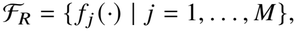

where *M* is the number of genes and each *f* _*j*_ (·) independently predicts the expression level of the gene *j* .

The patch image 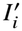 is processed by *f*_*PatchImg*_ (·). The resulting visual feature representations are then passed to the visual feature extractor of instances together with the proposals mask. *f*_*InstanceImg*_(·) extracts each proposal’s information by pooling with its proposal mask. The resulting visual feature representations of proposals and spot expressions are then passed to the group of regressors to predict each individual proposal’s gene expression. Since only spot-level gene expression *y*_*i*_ is available as supervision, instance-level predictions are aggregated by

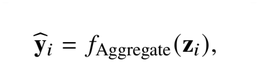

which sums instance-level outputs to yield a bag-level (spot-level) prediction. This formulation is consistent with the multiple-instance, multiple-target regression setting.

#### Training of regressors

The regressors module is trained under a weakly supervised learning framework, where the ground-truth labels are available only at the spot level. In this setting, the choice of loss function must ensure robustness to outliers and account for heteroscedastic variance across genes.

We therefore employ the Huber loss rather than the mean squared error (MSE) loss. The Huber loss between the predicted and ground-truth spot-level expressions is defined as

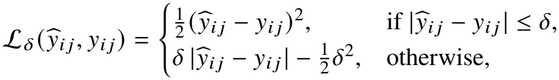

where *δ* is the threshold parameter that balances sensitivity and robustness. The total loss is averaged over all spots and genes as

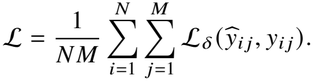

#### Implementation

To demonstrate that the proposed Vision Transformer–like architecture can be used effectively on relatively small spatial transcriptomics datasets, we employed a compact configuration: the Vision-Gene Transformer (VGT) consists of one encoder layer with three attention heads. Owing to the computational complexity, the number of genes used in training the VGT was selected based on the data. We used a size filter of 10 as the default, and applied h-maxima-based merging [11] as a post-processing step to remove noise objects and merge fragmented cells. For the regressors, we used the Adam optimizer [12] with a learning rate of 0.003. The visual feature extractor of the patch, *f*_*PatchImg*_ (·), used a group of convolutional layers (Supp. Fig. 1B); and each regressor *f*_*j*_(·) used a linear layer followed by a softplus activation unit [13]. To infer the count, we enforced integerization in the final stage of regressor, which can be used optionally.

### 2.3 Experiment design

In this section, we describe the generation of Visium-like simulated data, and preparation of the Visium HD and Visium Human IPF lung used for evaluation. We also describe methods used in these evaluations.

#### Algorithm 1

Attention Gradient Flow Tracking

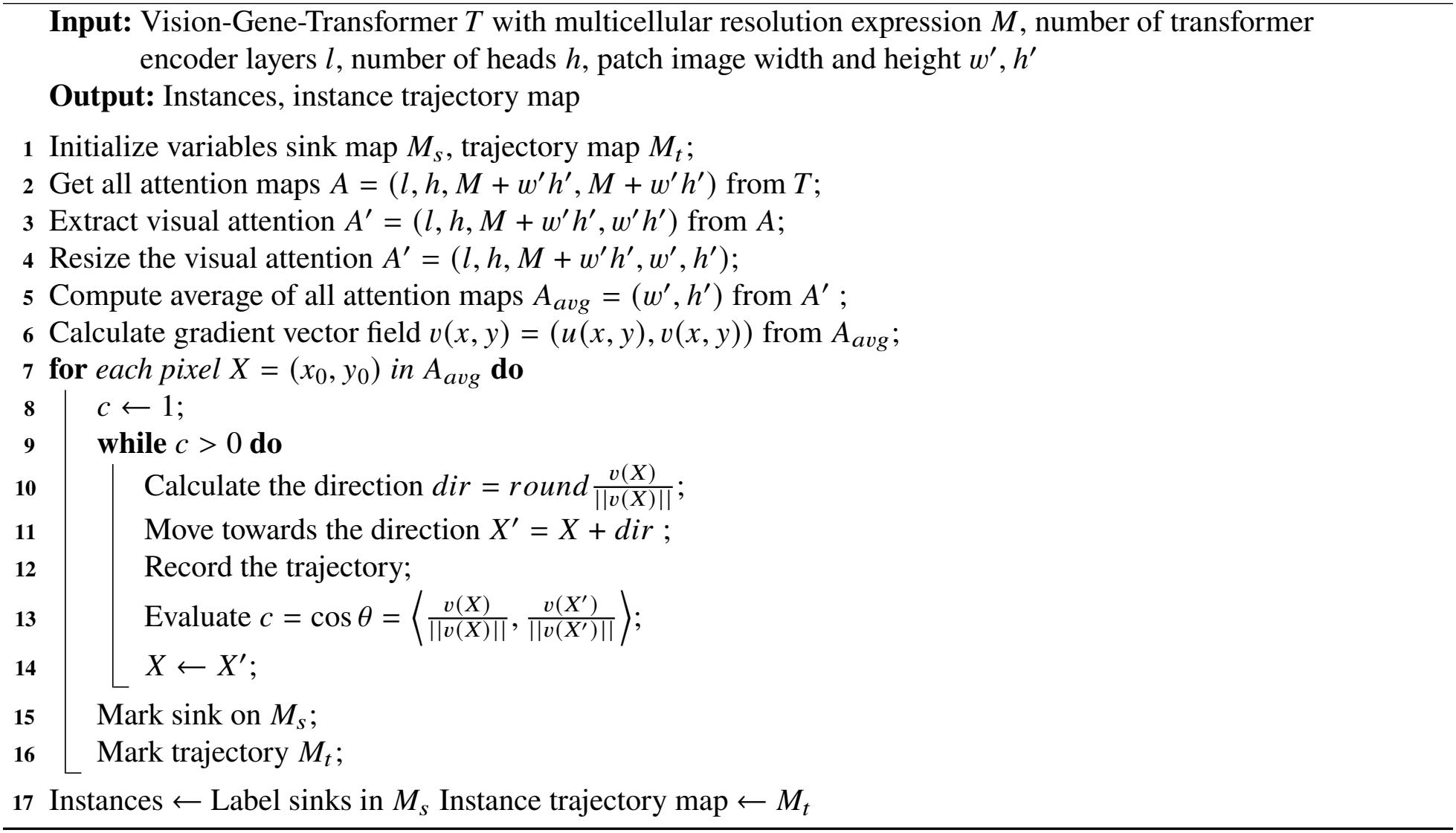

#### Visium-like simulated data

The 10x Xenium FFPE Human Breast dataset [14] was used to provide both the histology image and single-cell expression profiles for constructing Visium-like simulated data. From the histology image, we cropped 65 *µm* × 65 *µm* patches, with 100 *µm* between patch centers to mimic the Visium configuration. For each patch, cells located within the corresponding region were selected, and their aggregated gene expression profiles were recorded to represent spot-resolution spatial transcriptomics. The expression profile of each individual cell and its spatial area in the histology image were also recorded separately for evaluation. Patches containing no cells were excluded. In total, 3,971 spots with 313 genes were retained. We trained the st2sc VGT of *cell identification stage* using all 313 genes, with the global feature extractor processing 135 *µm* × 135 *µm* patches to provide additional spatial context.

We compared our method with two state-of-the-art approaches: SpatialScope [4] and scResolve [5], both of which infer single-cell-resolution spatial transcriptomics from multicellular spatial transcriptomics data. The input data were formatted as .h5ad files to meet the requirements of each method. scResolve was run with the configuration described in its original publication. SpatialScope, which requires a reference, used a breast cancer single-cell RNA-seq dataset [15] as its reference. Because of the high computational costs of both the construction of the reference model and gene expression decomposition step in SpatialScope, and to ensure a fair evaluation, 20,000 cells from 8 cell types (B-cells, CAFs, Cancer Epithelial, Endothelial, Myeloid, Normal Epithelial, PVL, T-cells) were selected from [15], and cell type annotations were restricted to this subsample. Cell type markers were defined as the top 20 genes by log-fold change for each cell type in the reference, and each predicted cell was assigned the cell type whose markers overlapped most with its 20 most highly expressed genes. To evaluate subtype marker genes, CD4+ T cells, CD8+ T cells, ACTA2+ myoepithelial and KRT15+ myoepithelial cells were selected from the annotation provided with the Xenium data. The *Cell identification stage* was evaluated by overlaying the center of each predicted cell on the ground-truth cell mask. Ground-truth cells were included in this evaluation if their centers lay within a spot. Gene expression was evaluated using all ground-truth cells in the spots: each ground-truth cell was compared with the predicted cells whose positions fell within its area. We measured three metrics of overall performance: (1) the mean squared error (MSE) between the expression of each ground-truth cell and its predicted cell; (2) cell type accuracy, comparing the cell types of the ground-truth and predicted cells; and (3) cell type correlation. For each spot, we correlated the cell type proportion vector of ground-truth cells with the cell type proportion vector constructed from each method. Cell identification and MSE were also compared at the cell type level. Finally, for subtype marker evaluation, we first determined, for ground-truth cells with known subtypes, whether the predicted cell type matched the ground-truth cell type. Only cells assigned their correct major cell type (T cells for the CD4+ and CD8+ subtypes, normal epithelial cells for the myoepithelial subtypes) were retained, so that this analysis measures subtype resolution independently of cell type accuracy. For each marker gene (CD8A, CD8B, CD4, KRT15 and ACTA2), we then computed the Spearman correlation between the predicted and measured counts across the retained cells. We used rank correlation because the methods report expression on different scales and marker counts are sparse and skewed (e.g., 32% of CD4+ T cells have no detected CD4 transcripts). Significance was assessed with a one-sided Mann–Whitney U test for MSE, a one-sided paired Wilcoxon test for cell type correlation, and a two-sided test of the Spearman correlation for subtype markers.

#### Visium HD

We used a mouse small intestine Visium HD dataset, which provides both the histology image and the 2 *µ*m expression bins. The data contain 19,059 genes and 5,479,660 2 *µ*m expression bins. st2sc can be applied to this near-single-cell-resolution technology. In the *cell identification stage*, we look at Visium-level image patches by aggregating the 2 *µ*m expression bins to a lower resolution. This allows cells to be identified largely as whole cells, rather than piece by piece from single bins, which would require an additional merging operation. In the *gene expression inference stage*, we train on 2 *µ*m expression bins. In the global feature extractor, we widen the view around each bin so that it covers the cells competing for that bin. When assigning the counts of a contested bin, the model looks at each identified cell as a whole, not only at part of the cell inside the bin, so that cell-level image information is used for accurate assignment.

The implementation details are as follows. We merged groups of 27 × 27 bins were merged to create 54 *µ*m × 54 *µ*m patches. Because of the computational cost of the large number of 2 *µ*m expression bins, we used 300 highly variable genes and 12,000 patches to train the *cell identification stage*, and then performed inference on all 13,381 patches. In the *gene expression inference stage*, 28 *µ*m image patches around each bin and all 19,059 genes were used. In this experiment, we evaluated our method against two methods widely used in Visium HD analysis, bin2cell[6] and ENACT[7]. Both assign the counts of 2 *µ*m expression bins to cells detected in the histology image. bin2cell was run with the configuration described in its tutorial, with count adjustment turned off so that raw counts were evaluated. For a fair comparison, ENACT was given the optional cell marker list, curated based on [16], and all other settings followed the original publication, including the “cluster weight” mode that splits shared bins. To annotate cell types, we used the snRNA-seq data of Drokhlyansky et al.[17], which contain 79,293 nuclei and 22,595 genes. We merged the subclusters into 19 major cell types and selected five marker genes per cell type from published studies to both annotate the cell types and curate the markers of cell types for the evaluation of assignment [17–20]. Each cell reported by each method was then assigned the cell type whose markers overlapped most with the cell’s 20 genes of highest relative expression among the 3,000 highly variable genes. Contested 2 *µ*m expression bins were not used for annotation, because they are used in the evaluation. Restricting the markers to ENACT’s gene set and removing markers with 5 or fewer counts in contested bins left 24 markers over 9 cell types.

#### Human Lung IPF Visium

A human idiopathic pulmonary fibrosis (IPF) lung Visium sample, together with human annotation, was obtained from the Cellular Senescence Network consortium (SenNet)[21]. The section comes from the lower lobe and was processed on the 10x Visium platform with Space Ranger. In total, 2,633 spots and 17,956 genes were measured. We used 55 µm × 55 µm spot image pathces. Training of VGT in *cell identification stage* in st2sc used 100 highly variable genes, with the global feature extractor observing 99 µm × 99 µm patches that cover the margin.

To build a reference for both cell type annotation and biological reference, we used Xenium IPF samples from [22]. 16 IPF samples were used. To train SpatialScope reference model and for cell type annotation of st2sc and scResolve, we used 7 cell types (Endothelial, Epithelial, Lymphoid, Macrophage, Fibroblasts, Multiciliated, Mesenchymal) including our cell type of interests (fibroblasts, macrophages) with 21,000 cells with 343 genes. The same cell type annotation method used in Visium-like simulated data was applied to st2sc and scResolve. To prepare gradient decay of fibroblast test, 8 out of 16 IPF samples annotated “Fibroblastic focus” label were used. Filtering one sample with less than 20 cells in fibrotic foci, and clustering cells into individual foci with 50 *µm* gap resulted in 20 fibrotic focus with 598 fibroblast cells. From each sample, we included fibroblast cells within 500 *µm* to measure the expression gradient distant from fibrotic focus. We computed the transcript fraction of fibroblasts in each 50 *µm* distance, then ordinary least-squares slope was used to get the gradient, Monte Carlo permutation test was used to measure the significance. The null distribution was generated from each focus to relocated to random position within its own sample then slope was recomputed. For SPP1+macrophage, macrophages annotated with at least one SPP1 expression count were relabeled as SPP1+macrophage.

## 3 Results

In this section, we compare cell positioning between the imaging-based cell identification method and attention gradient flow tracking against state-of-the-art approaches, as well as evaluate single-cell profiling from multicellular spatial transcriptomics methods in terms of absolute gene counts using Visium-like datasets. Next, we evaluate resolving the contested bin scenario in Visium HD against methods in Visium HD. Last, we show our application in human IPF lung Visium data to evalulate agreement with established biological findings.

### 3.1 st2sc recovers single-cell profile supported by Visium-like simulation

To show that st2sc recovers single-cell expression profiles from low-resolution spatial transcriptomics, we designed a Visium-like simulation using human breast cancer Xenium data (see Methods for details). In this evaluation, we compared st2sc with single-cell recovery methods that provide a full pipeline: SpatialScope, which relies on reference data to recover single-cell expression, and scResolve, which infers a super-resolution map of the expression profile. Since all pipelines include both a *cell identification stage* and a *gene expression inference stage*, we first evaluated *cell identification stage*. Because scResolve and SpatialScope report only cell center coordinates rather than full cell areas, we evaluated the *cell identification stage* of all methods by measuring whether each predicted center of the cell position overlays a ground-truth cell. st2sc identified the most cells (49,242 cells, 69.5% of the cells whose centers lie within a spot), whereas SpatialScope (39,326, 55.5%) and scResolve (19,687, 27.8%) identified fewer cells (Fig. 3A). We further examined performance at the cell type level (Supp. Fig. 2A). Across all cell types, st2sc identified more cells than the other methods, indicating that its performance is robust across cell types.

**Fig. 3.**
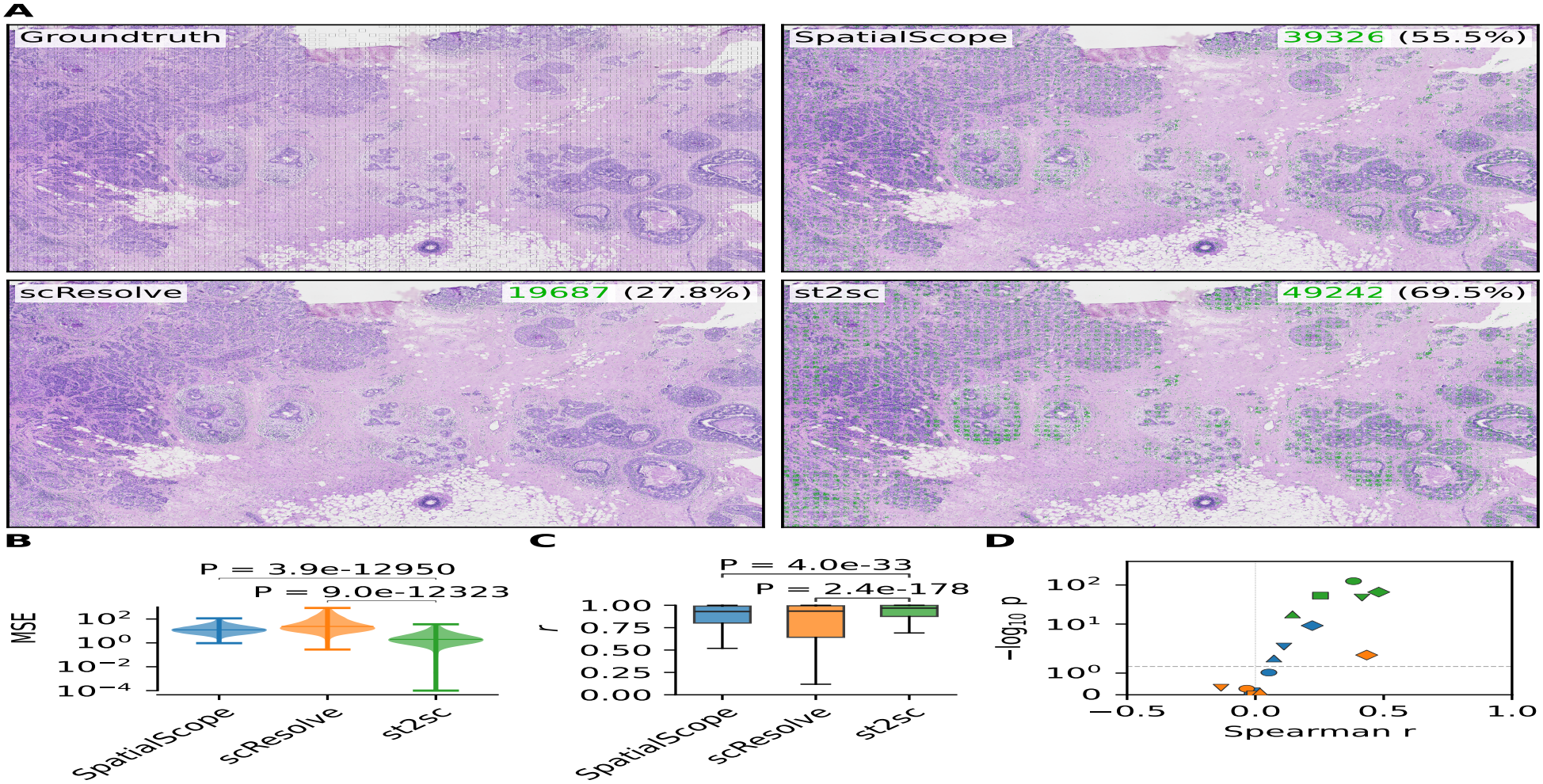
st2sc recovers single-cell expression in a Visium-like simulation. (A) ground-truth histololgy image and evaluation of *cell identification stage*. Grid box shows area used to simulate low resolution simulation. Green dots show predicted cell overlaid on the ground-truth cell. (B) Per-cell mean squared error (MSE) between ground-truth and predicted expression. Lines mark the median (C) Per-spot Pearson correlation coefficient (PCC) between ground-truth and predicted cell type proportions on the spots covered by all methods. (D) Recovery of subtype marker expression. Spearman correlation between ground-truth and predicted marker counts. Dashed line denotes *p* = 0.05. Circle: CD8+ T cell - CD8A. Square: CD8+ T cell - CD8B. Triangle: CD4+ T cell - CD4. Diamond: KRT15+ Myoepi cell - KRT15. Reverse triangle: ACTA2+ Myoepi cell - ACTA2

Next, we evaluated the *gene expression inference stage*, measuring how accurately st2sc recovers single-cell gene counts by mean squared error (MSE), and compared it with SpatialScope and scResolve (Fig. 3B). st2sc achieved an MSE of 3.6833, significantly outperforming the other methods (SpatialScope: 18.2552, *p* = 3.9e ™ 12950; scResolve: 65.3402, *p* = 9.0e ™ 12323). At the cell type level, st2sc also outperformed the other methods across all cell types, with MSE 3.9-fold (CAFs) to 16.3-fold (T cells) lower than SpatialScope, indicating that this robustness extends to gene expression inference (Supp. Fig. 2B). Both st2sc and scResolve benefit from not relying on a reference, inferring all 313 genes, whereas SpatialScope is limited to the 308 genes shared with its reference.

We further investigated performance in two ways. First, if the expression profile is correctly recovered, the cell type should also be inferable. To evaluate this, we measured both cell type accuracy and per-spot cell type correlation. st2sc achieved 69.50% and 0.8911, respectively, whereas lower performance was observed for both SpatialScope (52.77%, 0.8584) and scResolve (65.72%, 0.7534) (Fig. 3.C, Supp. Fig. 2C). Second, to assess recovery beyond overall performance, we tested whether our method recovers expression patterns of subtype-specific markers (see Methods for details). st2sc achieved the highest Spearman correlation for subtype markers (0.481 for KRT15 in KRT15+ myoepithelial cells; 0.382 for CD8A in CD8+ T cells), whereas SpatialScope (0.222, 0.053) and scResolve (0.434, ™0.034) showed lower correlations, indicating weaker recovery of subtype marker expression (Fig. 3D). Together, these results show that recovering single-cell profiles from spot counts is possible with st2sc, and that it does so accurately enough to capture within-type variation.

### 3.2 st2sc resolves the contested bin in Visium HD

To test whether st2sc assigns expression counts to the cells they belong to in Visium HD, we evaluated the methods on 2 *µ*m expression bins that are “contested” by two cells. Because ground-truth expression count assignments are unavailable, we evaluated assignment quality using cell-type-specific marker genes as a proxy. Under the assumption that these markers are preferentially expressed by their corresponding cell types, higher fractions of marker counts assgined to the expected cell types indicate more biologically consistent assignment. This naturally restricts the evaluation to bins contested by two cells of different cell types (Fig. 4A). This analysis also allowed us to assess to show that st2sc can be applied to a different modality.

**Fig. 4.**
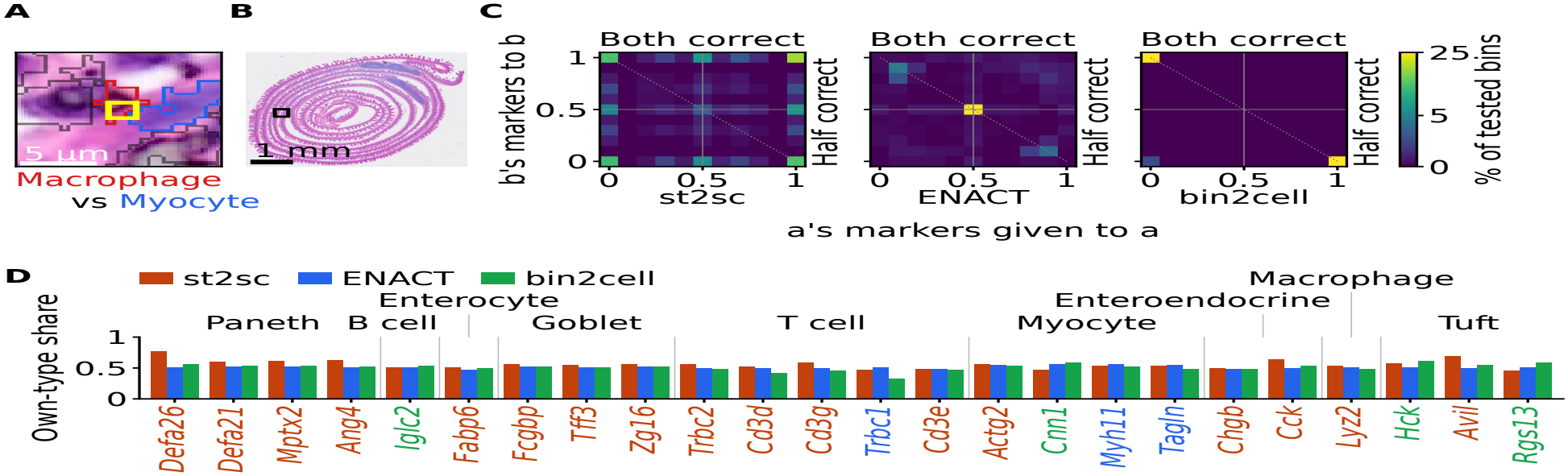
st2sc assigns counts in the contested bin in Visium HD. (A) Example image of marcophage and myocyte overlapped in a 2 *µ*m expression bin (B) Mouse small intestine Visium HD used in this study. (C) 2D histogram showing assignment performance for each method. (D) Bar plot showing gene level assignment assessment for each method.

st2sc can incorporate a larger image context with the global feature extractor module. Based on this ability, once cells are identified in *cell identification stage* and contested bins are located, we first trained *gene expression inference stage* using bins overlapping a single identified cell, where we sum the expression covered by the cell. Then with tested bins, we assigned marker gene counts for each cell.

We used the Visium HD mouse small intestine data to establish the evaluation pipeline (Fig. 4B; see Methods for details). We first built a 2D histogram for the evaluation (Fig. 4C). When only one of the two cells receives its corresponding marker counts, the bin falls in the bottom-right or top-left cell of the histogram. When both cells receive their corresponding marker counts, the bin falls in the top-right cell. st2sc reported 4,787 bins contained two cells of different cell types with markers of both types (tested bins) and in 18.6% of them (889 bins) both cells received all of their corresponding marker counts. In contrast, ENACT reported 14,744 tested bins, of which only 1 bin had both cells received at least 95% of their corresponding marker counts, and bin2cell reported 4,076 tested bins with no bins in the top-right cell. Even when we expanded the analysis to the whole top-right quadrant, st2sc performed best (1,123 bins, 23.5%), compared with ENACT (325 bins, 2.2%) and bin2cell (0 bins). bin2cell cannot reach this quadrant because it assigns all expression counts of a bin to one cell, which produces bright cells at the top-left and bottom-right of the histogram. Unlike the baseline methods, st2sc produced a substantial number of tested bins in which marker counts for both cell types were predominantly assigned to their corresponding cells, suggesting an advantage in resolving overlapping cellular contribtuions. We then moved from the overall result to individual cell types and marker genes (Fig. 4D). st2sc assigned markers better in 6 of 9 cell types, while bin2cell performed best for two cell types (B cell and Tuft) and ENACT won in Myocyte. st2sc achived its highest assignment percentage for Defa26: 52 of 68 UMIs (76.5%) in tested bins were assigned to the Paneth cell, whereas ENACT (137 of 269, 51.0%) and bin2cell (37 of 66, 56.1%) assigned it less well. Defa21 accounted for the largest number of the UMIs in Paneth counts, and st2sc showed 2873 of 4832 (59.5%) while ENACT (9606 of 18581, 51.7%) and bin2cell (2,444 of 4,522, 54.0%) showed less assignment. Across the four Paneth marker genes, st2sc assigned 4,343 of 7,173 UMIs (60.5%) to the corresponding cell, while ENACT (14,213 of 27,521, 51.6%) and bin2cell (3,563 of 6,640, 53.7%) showed less assignment. Another cell type where st2sc showed robust assignment was Goblet, where st2sc assigned 12,579 of 22,495 (55.9%) to the corresponding cell across three marker genes, while ENACT (35,793 of 68,988, 51.9%) and bin2cell (11,447 of 22,048, 51.9%) again showed lower performance than st2sc. Interestingly, in most cell type st2sc performed best, st2sc achieved higher assignment fractions for all evaluated marker genes of that cell type. This shows that the advantage of st2sc’s *gene expression inference stage* is consistent across the markers of a type, not driven by single genes. These results show that st2sc improves the assignment of contested bins in Visium HD.

### 3.3 st2sc recovers expression gradient decay of fibrotic foci and SPP1+macrophage accumulation

We next investigated whether single-cell expression profiles inferred from low-resolution spatial transcriptomics data could capture spatial patterns previously observed using higher-resolution spatial transcriptomics. We focused on two biological features associated with idiopathic pulmonary fibrosis (IPF): changes in fibroblast gene expression with distance from fibrotic foci and the spatial organization of SPP1+ macrophages. For these evaluations, we selected a human IPF lower-lobe Visium sample with expert annotations of fibrotic foci (Fig. 5A) from the Cellular Senescence Network [21]. Because spot-level expression represents aggregated signals from multiple cells, we hypothesized that recovering individual cellular expression profiles could enable the identification of cell-type-specific spatial patterns that are not directly accessible from the low resolution measurements. To establish a reference pattern for the first analysis, we examined Xenium IPF lung samples with expert-annotated fibrotic foci [22]. Using these annotations, we investigated whether fibrosis-associated genes exhibited expression gradients in fibroblasts as a function of distance from the fibrotic foci. We evaluated five genes (COL1A1, COL1A2, POSTN, CTHRC1, and TGFB1) based on previous studies [23–26]. Among these genes, three (COL1A1, CTHRC1, and TGFB1) exhibited significantly decreasing expression in fibroblasts with increasing distance from the fibrotic foci (Fig. 5B).

**Fig. 5.**
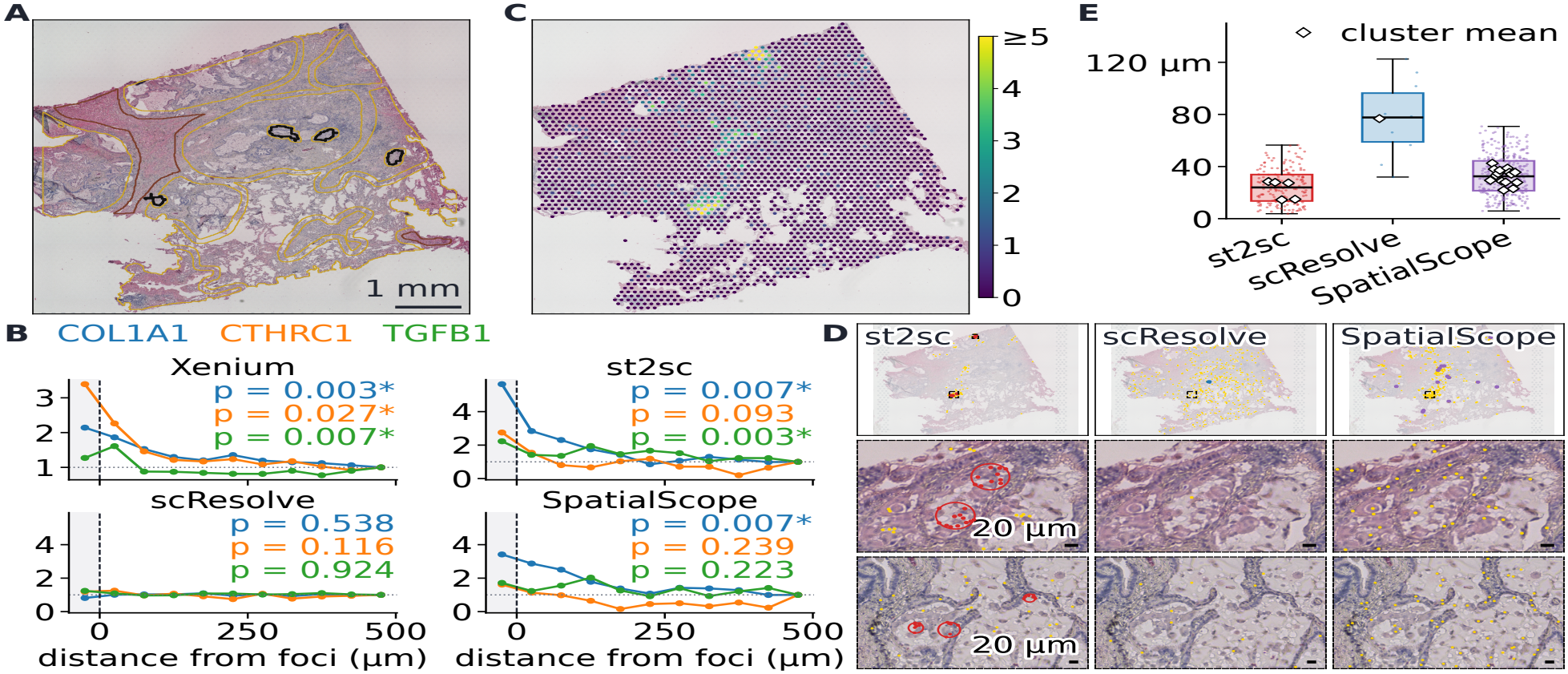
st2sc recovers spatial patterns consistent with known IPF biology from Visium. (A) IPF lower lobe sample with expert annotation used in this study. Black line: Fibrotic foci Yellow line: Bronchiolo-vascular. Red line: Old hyalinised scar. (B) Expression gradient decay of fibroblasts from fibrotic foci. Y axis denotes transcript fraction relative to 450-500 *µ*m. Blue: COL1A1 Orange: CTHRC1. Green: TGFB1. (C) SPP1 count in spot resolution. (D) SPP1+macrophages accumulation in airspace. (E) Distance between SPP1+ macrophages and within-cluster pair distance. Each dot represents pair distance between SPP1+ macrophages. Diamond represents average distance of SPP1+macrophages in identified cluster. Y axis denotes within cluster distance (*µ*m)

We then evaluated whether single-cell expression profiles inferred from Visium data could reproduce these fibroblast-specific expression gradients. st2sc recovered significant decreasing expression gradients for COL1A1 and TGFB1, whereas SpatialScope recovered a significant gradient only for COL1A1, and scResolve did not recover significant gradients for any of the three genes (Fig. 5B). These results suggest that st2sc can recover cell-type-specific spatial expression patterns consistent with those observed using higher-resolution spatial transcriptomics, despite the limited spatial resolution of conventional Visium measurements.

Having evaluated fibroblast-specific expression gradients, we next investigated whether st2sc could recover the spatial accumulation of SPP1+ macrophages, another biological feature associated with pulmonary fibrosis. *Vannan et al*. reported accumulation of the SPP1+macrophage in the airspace is observed as progression of the pulmonary fibrosis [22]. Based on the existence of multiple fibrotic foci and old hyalinised scar annotation, we considered this sample has enough progression of the pulmonary fibrosis to show accumulation of the SPP1+macrophage. SPP1 count in the visium showed enriched in three distinct regions, and most of the spots outside of these three regions had count of 1 (Fig. 5C). When we annotated SPP1+macrophage using SPP1 expression, scResolve and SpatialScope showed overidentified the SPP1+macrophage in those count of 1 region (Fig. 5D), reporting wide but sparse clusters. On the other hand, st2sc reported SPP1+ macrophages focused on these 3 regions, and showed more densely populated SPP1+macrophages forming spatial clusters, which aligns with the report. The results of these applications in human IPF lung show st2sc reveals biological findings reported in higher resolution spatial transcriptomics from low resolution spatial transcriptomics.

## 4 Discussion

Existing computational methods aimed at inferring single-cell profiles from multicellular spatial transcriptomics have emphasized the prediction of cell type as the primary performance metric. However, this focus often overlooks the need for accurate gene count prediction to achieve accurate profiling. To address this gap, we propose the st2sc framework, specifically designed to infer true single-cell gene expression within a multiple-instance, multiple-target regression problem. In particular, we introduced a novel attention gradient flow tracking algorithm to instantiate cells from an input histology image, as well as provided a deep learning-based approach to predict single-cell profiles. Beyond improving overall cell-type identification, the results highlight the importance of recovering gene-level variation within individual cells. In the Visium-like simulation, improved expression inference enabled the recovery of selected subtype-marker patterns that may not be adequately characterized by cell-type annotations alone. In the IPF lung analysis, inferred cell-level profiles captured spatially organized gene-expression patterns associated with fibroblasts and macrophages. These findings suggest that cell-resolved expression inference may provide a complementary analytical perspective to spot-level deconvolution, particularly when spatial variation within a cell type is biologically informative.

While we proposed a way to remove unwanted objects in the postprocessing of attention gradient flow tracking, this does not completely remove spurious instances or merge all fragmented instances. We believe that there can be further postprocessing using the angle map, as each pixel is bounded between -1 and 1. Furthermore, the approach could further developed for the accurate segmentation of the cell, as currently the model shows potential of capturing the boundary based on both histology and expression from VGT. An interesting future direction involving the use of attention gradient flow tracking is the quantitative analysis of the instances using attention-derived probability. If the instance mask is produced over the probability map, the summation of the probability pixels provides a direct probabilistic score for each instance. This probability for each object in certain head can be used for quantitative evaluation of the multi-head, which could provide a principled approach for selecting informative attention heads in the multi-head transformer architecture.

## 5 Acknowledgement

This work was partially supported by National Institutes of Health (NIH) grant 1U54AG075931 and National Science Foundation (NSF) award 2134998 (to J.L.-M.).

