## Supplementary document for "Inferring single-cell gene expression profiles from multicellular spatial transcriptomics data with st2sc"

### Appendix

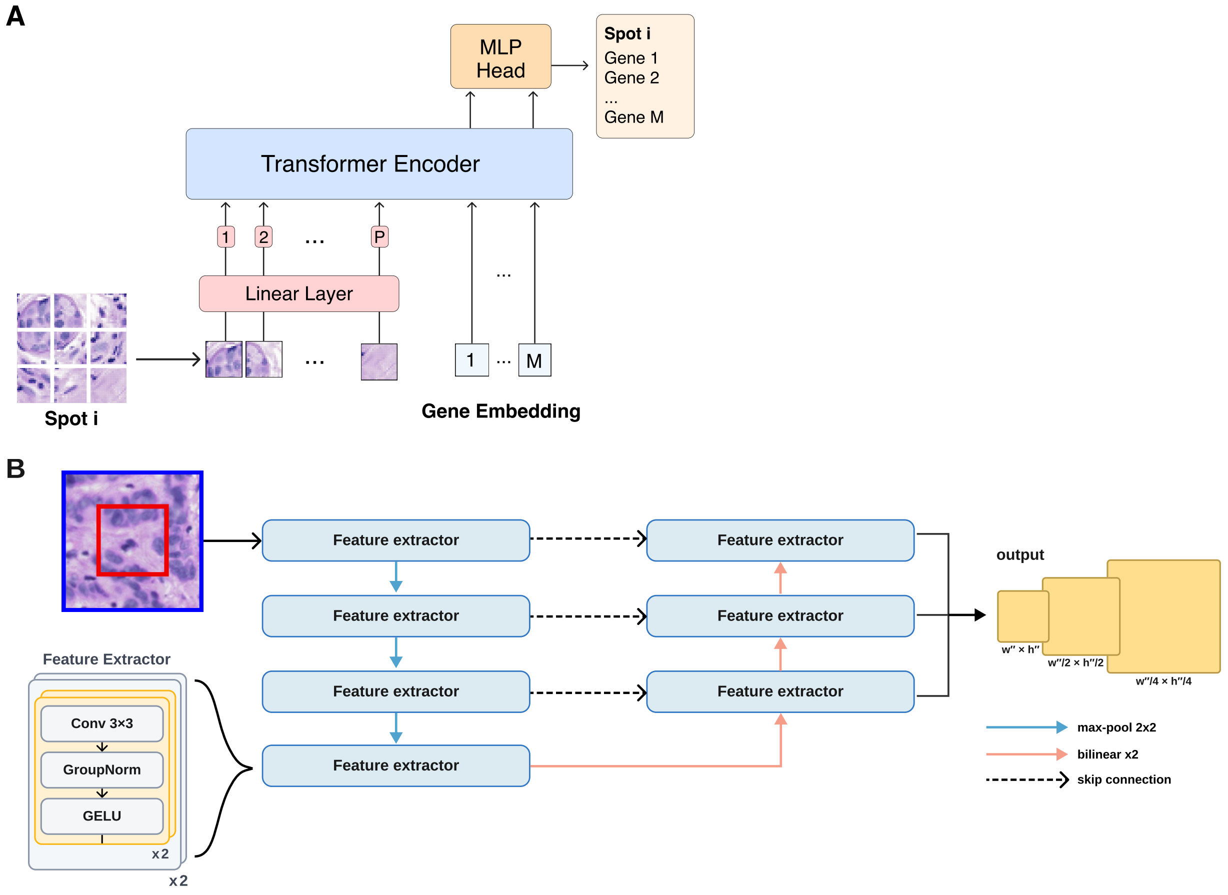


**Supp. Fig. 1. Model components in st2sc** (A) Architecture of the Vision-Gene Transformer (VGT). In this model, the class embedding of a standard Vision Transformer is replaced with a gene embedding, enabling the network to attend to cellular regions relevant for inferring spot-level gene expression. (B) Architecture of the Global feature extractor. The patch image is used in this model to provide image of the cells trimmed in the spot image and more context of the nearby tissue environment.

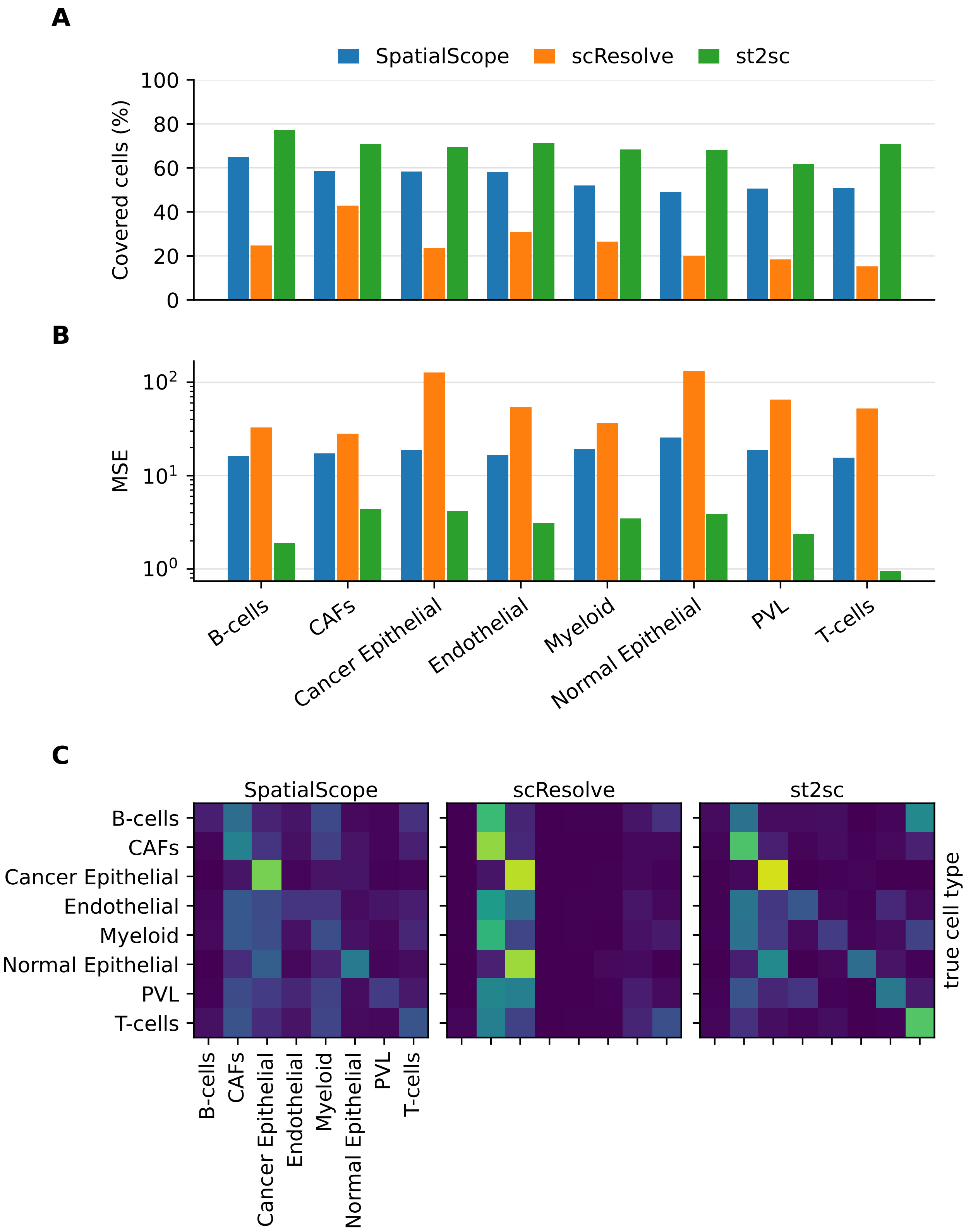


**Supp. Fig. 2. Extended figure of Visium-like simulation** (A) *Cell identification stage* performance by cell type. (B) Mean squared error (MSE) of gene expression inference by cell type in a log scale. (C) Confusion matrices of cell type assignment for each method. Rows are ground-truth cell types. Columns are predicted cell types.

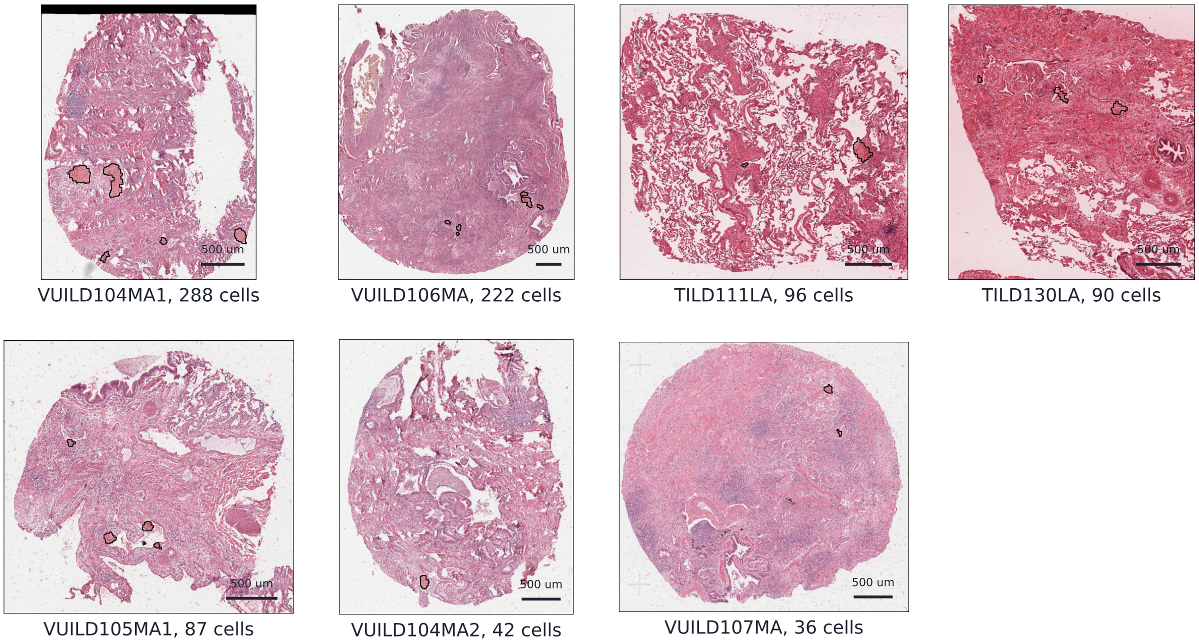


**Supp. Fig. 3. H&E images and annotation of the seven IPF samples.** Samples from Vannan *et al.* in which cells carry the pathologist label "Fibroblastic Focus". Black outlines enclose the labelled cells. Each panel is labelled with the sample name and its total cell count.
